# Imaging sensory transmission and neuronal plasticity in primary sensory neurons with genetically-encoded voltage indicator, ASAP4.4-Kv

**DOI:** 10.1101/2021.05.21.445202

**Authors:** Yan Zhang, John Shannonhouse, Ruben Gomez, Hyeonwi Son, Hirotake Ishida, Stephen Evans, Mariya Chavarha, Dongqing Shi, Guofeng Zhang, Michael Z Lin, Yu Shin Kim

## Abstract

Detection of somatosensory inputs requires conversion of external stimuli into electrical signals by activation of primary sensory neurons. The mechanisms by which heterogeneous primary sensory neurons encode different somatosensory inputs remains unclear. *In vivo* dorsal root ganglia (DRG) imaging using genetically-encoded Ca^2+^ indicators (GECIs) is currently the best technique for this purpose by providing an unprecedented spatial and populational resolution. It permits the simultaneous imaging of >1800 neurons/DRG in live mice. However, this approach is not ideal given that Ca^2+^ is a second messenger and has inherently slow response kinetics. In contrast, genetically-encoded voltage indicators (GEVIs) have the potential to track voltage changes in multiple neurons in real time but often lack the brightness and dynamic range required for *in vivo* use. Here, we used soma-targeted ASAP4.4-Kv, a novel GEVI, to dissect the temporal dynamics of noxious and non-noxious neuronal signals during mechanical, thermal, or chemical stimulation in DRG of live mice. ASAP4.4-Kv is sufficiently bright and fast enough to optically characterize individual neuron coding dynamics. Notably, using ASAP4.4-Kv, we uncovered cell-to-cell electrical synchronization between adjacent DRG neurons and robust dynamic transformations in sensory coding following tissue injury. Finally, we found that a combination of GEVI and GECI imaging empowered *in vivo* optical studies of sensory signal processing and integration mechanisms with optimal spatiotemporal analysis.

**Highlights:**

- *In vivo* ultra fast and sensitive dynamic voltage imaging of peripheral primary sensory neurons by a newly generated genetically-encoded voltage indicator.
- Identification of mechanical, thermal, or chemical stimuli-evoked voltage signals with superior temporal resolution.
- Single-cell detection of changes in sub- and suprathreshold voltage dynamics across different disease conditions.
- Combination of voltage (by ASAP4.4-Kv) and Ca^2+^ (by Pirt-GCaMP3) signals to facilitate the understanding of signal processing and integration of primary sensory neurons, especially for noxious versus non-noxious sensation.

## Introduction

Dorsal root ganglia (DRG) neurons have pseudounipolar axons that project toward skin where they initially convert external stimuli such as touch, stretch, itch, hot, cold, and/or chemical stimuli into corresponding electrical signals. These electrical signals are integrated and modulated in the cell bodies of DRG located in intervertebral foramen between spinal vertebrae, and then action potentials containing somatosensory information are further propagated to the superficial laminae of dorsal spinal cord. Electrophysiologic recording has been used as a fundamental tool for measuring neuronal electrical signals for many decades, but this approach is limited by the invasiveness of the procedure, poor anatomical accessibility, the absence of physiological input during commonly used *in vitro* recordings, and by the difficulty of *in vivo* measurement due to stability issues^1–3^. Genetically-encoded Ca^2+^ indicators (GECIs) allow for monitoring DRG neuronal firing activities, network patterns among neurons and other cell types, and sensory circuits in physiological and pathological conditions with exceptional spatial and populational resolution with limited perturbation^4^. However, Ca^2+^ indicators fail to distinguish between action potential-evoked Ca^2+^ influx vs. Ca^2+^ transients arising from internal stores and ligand-gated Ca^2+^ channels. Furthermore, Ca^2+^ indicators only report suprathreshold signaling while failing to detect subthreshold membrane potential fluctuations due to slow kinetics and limited sensitivity^2,3,5^. As an alternative, recording DRG neuronal activity using fluorescence generated from genetically-encoded voltage indicators (GEVIs), which can follow not only fast suprathreshold voltage signals but also subthreshold fluctuations in membrane potentials, could be an excellent complementary approach to *in vivo* GECI imaging.

Dynamic voltage imaging employing voltage sensing dyes and later with GEVIs has been used for decades to study electrical activity in various tissues and organisms *in vitro* and *in vivo*^6–10^, yet no data are available pertaining to functional voltage imaging of primary sensory neurons due to the lack of appropriate tools and techniques. More recently, GEVIs have been developed that enable stable expression in mammalian cells^11^, allowing the use of genetic tools to achieve cell type specificity^2,3^. Despite the widespread use of GEVIs, only a handful have been successfully used for *in vivo* optical imaging to detect voltage dynamics in a living mouse brain. These include archaerhodopsin-based indicator Ace2N, paQuasAr3-s, and SomArchon^12,13,14^, and ASAP-family GEVIs in which a voltage-sensitive domain (VSD) is linked to a circularly permuted GFP protein^15–17^. Compared to archaerhodopsin-based GEVIs, ASAP3-Kv enables accurately tracking suprathreshold and subthreshold voltage dynamics with decent signal-to-noise ratio (SNR) from deep brain regions in live mice^15,16^. However, ASAP3-Kv generates a reversed optical signal (a decrease in fluorescence intensity with membrane depolarization), which leads to high excitability and thus high photobleaching at resting membrane potentials. Existing positively tuned GEVIs that brighten with positive voltage changes use either VSDs from voltage sensitive phosphatases (ElectricPk^18^, FlicR1^19,20^ and Marina^21^), or electrochromic fluorescence resonance energy transfer (eFRET) like Ace2N-mNeon^22^. Among these indicators, Marina exhibits largest optical responses in *in vitro* experiments while practical use *in vivo* needs more evaluations. Recent attempts have successfully developed some positively tuned ASAP4 voltage indicators that surpass currently available positively tuned voltage indicators in their fluorescence responses and SNRs^17^. One novel ASAP4-subfamily GEVI, ASAP4.4-Kv, which ASAP4.4 voltage indicator is attached to the Kv2.1 potassium channel to locate ASAP4.4 to the soma, combines notable properties for *in vivo* applications: brighten in response to membrane depolarization, high-level neuronal expression, fast kinetics, and large fluorescence changes. We therefore anticipate that the properties of ASAP4.4-Kv make it more suitable and optimal for routine and robust *in vivo* DRG imaging by reporting action potentials reliably and revealing subthreshold events in optical studies.

To illustrate the feasibility and utility of using ASAP4.4-Kv as an indicator in primary sensory neurons, we examined neuronal activity in the DRG of live mice. Data acquired using ASAP4.4-Kv uncovered striking cell-to-cell communication and synchronization patterns between adjacent DRG neurons following peripheral inflammation or nerve injury, which was rarely seen in naïve animals. ASAP4.4-Kv data permitted visualizing the representation of mechanical, thermal, and chemical stimuli *in vivo*, and enabled us to track down how these parameters transform with peripheral injury. By comparing *in vivo* GEVI and GECI signals, we found that GECI imaging can represent complex phenomena extending to an entire population ensemble of DRG neurons, but that images lack temporal precision and fidelity. We also found that ASAP4.4-Kv voltage imaging enabled visualization of temporal dynamics of individual DRG neurons with fast temporal resolution. We conclude that combining GEVI and GECI imaging provides an optimal approach for analyzing the complex signal processing and integration of somatosensation in various contexts.

## Results

### ASAP4.4-Kv imaging reveals electrical coupling synchronization between adjacent DRG neurons

For *in vivo* DRG voltage imaging, we intrathecally injected adeno-associated viruses (AAVs) encoding ASAP4.4-Kv into spinal cord to allow for expression in DRG neurons. At 5–7 weeks after injection, *in vivo* single photon confocal imaging experiments were performed on the right lumbar (L5) DRG, which innervates parts of right hindpaw, leg, and back of the mouse. Fluorescent signals from ASAP4.4-Kv were acquired by confocal microscopy in frame mode to capture the entire population of L5 DRG neurons (supplementary Fig. 1). We verified transduction of ASAP4.4-Kv virus into DRG neurons by imaging ASAP4.4 fluorescence in DRG neurons (Fig. 1a and supplementary Fig. 1). The basal ASAP4.4-Kv green fluorescence intensity was relatively low under *in vivo* conditions; however, inflammation in hindpaw caused by complete Freund’s adjuvant (CFA) injection or chronic constriction injury^23^ of sciatic nerves (SNs) yielded a stronger ASAP4.4-Kv fluorescent signal (Fig. 1a and supplementary Fig. 1). The results showed that ASAP4.4-Kv can be sparsely but highly expressed in DRG neurons *in vivo*, and can dynamically respond to voltage in the physiological range (supplementary Fig. 2). These are essential properties for carrying out the functional analysis at the cellular level *in vivo*.

**Figure 1.**
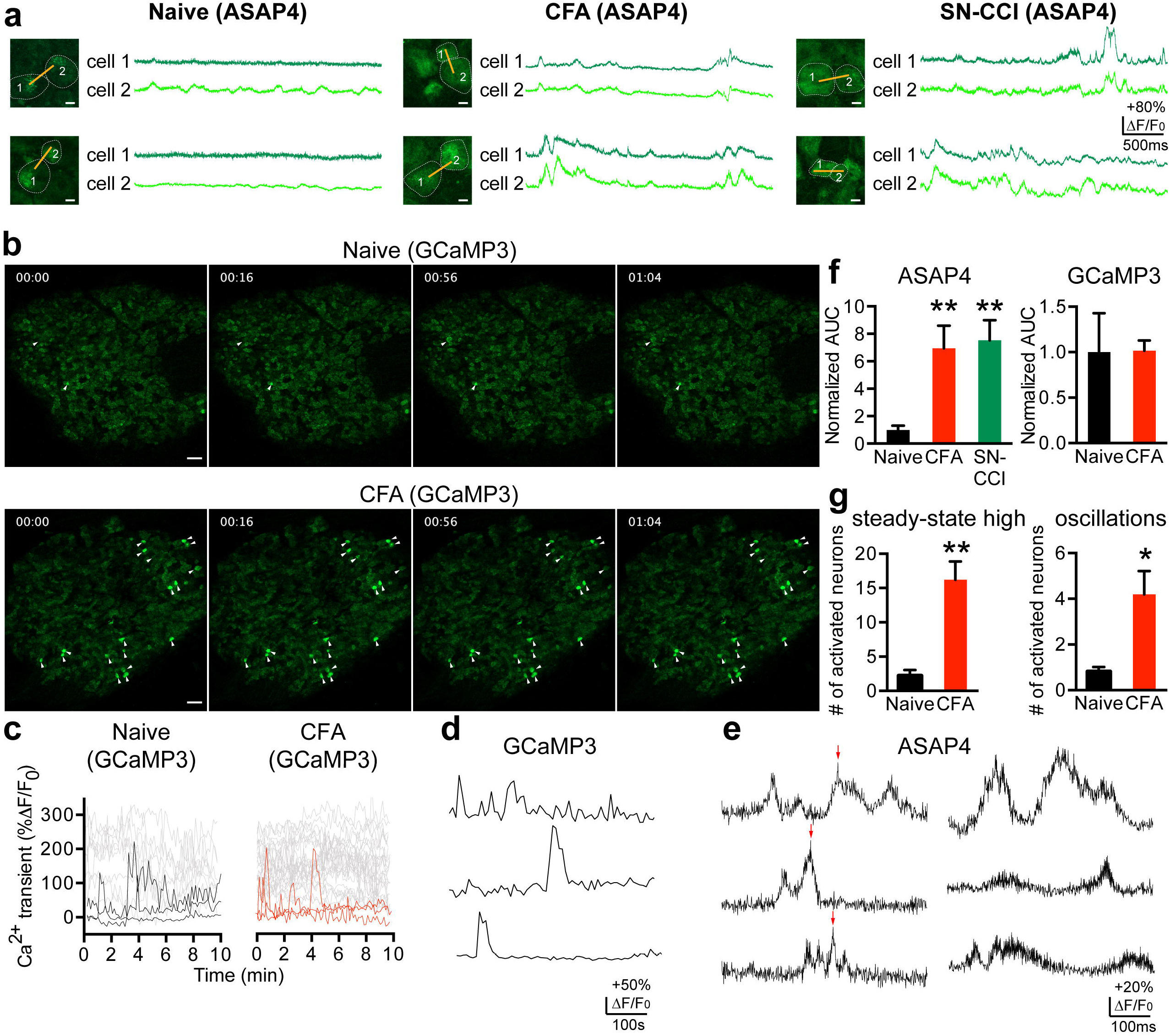
*In vivo* optical detection of electrically synchronous, spontaneous neuronal activity using ASAP4.4 voltage imaging in intact DRG neurons. **(a)** Images show two DRG somata expressing ASAP4.4. Yellow lines indicate 1.1 kHz line scan regions where ASAP4.4 fluorescent signals were acquired. Traces show simultaneous optical imaging of spontaneous electrical activity in two adjacent DRG neurons using the ASAP4.4 voltage sensor. Scale bar: 10 μm. **(b)** Spontaneous Ca^2+^ activity in DRG neurons measured optically with genetically-encoded Ca^2+^ indicator using Pirt-GCaMP3 mice. White arrowheads indicate spontaneously activated neurons in the absence of stimulus. Scale bar: 100 μm. **(c)** GCaMP3 reports spontaneously activated neurons with steady-state high Ca^2+^ levels (grey) and spontaneously activated neurons with Ca^2+^ oscillations (black or red) in the DRG (representative of three experiments for each treatment). **(d, e)** Optical ASAP4.4 **(e)** or GCaMP3 **(c, d)** traces show high temporal signal fidelity in ASAP4.4 voltage imaging (milliseconds) vs. GCaMP3 imaging (seconds to minutes). Red arrows indicate action potential at the peak of oscillation. **(f)** Mean area under the curve (AUC) of ASAP4.4 signals in L5 DRG neurons from naïve, CFA, or SN-CCI animals (n=3 mice/group; **P<0.01, Kruskal-Wallis test), and mean AUC of spontaneous Ca^2+^ oscillations in L5 DRG neurons from naïve or CFA-treated Pirt-GCaMP3 mice (n=3 mice/group; P=0.4643, Mann-Whitney U-test). **(g)** Total number of spontaneously activated neurons in L5 DRG from naïve or CFA-treated Pirt-GCaMP3 mice (n=3 mice/group; *P<0.05, **P<0.01, Mann-Whitney U-test).

Subthreshold voltage changes in membrane potential are mainly driven by the spatiotemporal integration and summation of somatosensory inputs. Ectopic discharge and subthreshold membrane potential oscillation in DRG neurons are underlying cellular mechanisms for pain behaviors in many injured animals^24,25^. Thus, we investigated the ability of ASAP4.4-Kv to detect spontaneous DRG neuronal activity *in vivo*. Both bright and dim cells in naïve or injured mice were chosen for line scan imaging for comparison. ASAP4.4-Kv voltage imaging was focused on superficial regions of DRG neuronal cell membranes at depths below the meninges ranging from 0 to 20 μm, and confocal imaging was performed in line scan mode at 1–1.4 kHz. Using ASAP4.4-Kv, we rarely detected subthreshold electrical signal changes in DRG somata of naïve mice without stimulus (Fig. 1a). Interestingly, subthreshold voltage fluctuations were readily detectable *in vivo* in the context of inflammation or nerve injury (Fig. 1a), with approximately 7-fold increase in average area under the curve (AUC) of ASAP4.4-Kv fluorescent signal intensity (Fig. 1f). These findings demonstrate the capability of ASAP4.4-Kv to uncover subthreshold voltage fluctuations in DRG neurons *in vivo*, which were previously inaccessible due to limitations of low-yield intracellular or whole-cell patch clamp electrophysiologic recording techniques. Close examination of the ASAP4.4-Kv signal trajectories revealed that the membrane potential changes were actually composed of periodically appearing subthreshold voltage oscillations and minor spikes which usually occurred from the peak of oscillation and lasted only a few milliseconds (Fig. 1e).

To detect neuronal cell-to-cell communication in the neuronal network and circuits in DRG, we randomly selected pairs of adjacent DRG neurons in different regions of the DRG, and performed a single-line scan at about 1.1 kHz across the membrane regions of the two neuronal cells (Fig. 1a, images). We analyzed paired data sets from naïve, CFA, or SN-CCI animals by quantification of fluorescence intensity changes in scanned areas of individual adjacent cells. In naïve mice, no temporal cell-to-cell coherence or synchronization of subthreshold voltage fluctuations in neuronal membranes was detected (Fig. 1a, left). Strikingly, under the context of inflammation or nerve injury, around 6% of recording neuronal pairs exhibited strong coincident subthreshold voltage changes in the range of ten to hundreds of milliseconds, regardless of activity patterns. (Fig. 1a, middle and right), whereas gap junction blocker, carbenoxolone (CBX), significantly reduced cell-to-cell electrical synchronization (supplementary Fig. 3). Our results indicate that tissue injury increased cell-to-cell connectivity and network communication between DRG neurons leading to enhanced synchronization in DRG neuronal networks, and eventually to better integration and summation of somatosensory signals. To the best of our knowledge, such electrically synchronous neuronal events between cells in the peripheral sensory system *in vivo* have not been previously described.

To determine whether electrically synchronous events corresponded to global neuronal activity, we included *in vivo* DRG Ca^2+^ imaging of neuronal populations using Pirt-GCaMP3 mice, in which the GECI GCaMP3 was exclusively expressed in primary sensory neurons under the control of the *Pirt* promoter^26^. Using Pirt-GCaMP3 Ca^2+^ imaging, we could simultaneously monitor neuronal activity of the entire population of DRG neurons^4^. We imaged the entire DRG at ~6.4 to 7.9 s/frame and found that spontaneous activity was rarely seen in naïve animals (1–3 neurons/DRG), but in the presence of inflammation or nerve injury, increased spontaneous neuronal activity was observed (>10 neurons/DRG) (Fig, 1b, c, d). This Ca^2+^ activity could represent either sporadic Ca^2+^ oscillations or steady-state high Ca^2+^. To this point, however, no synchronized spontaneous activity was observed in GCaMP3 signals. In comparing voltage dynamics seen by ASAP4.4-Kv imaging with Ca^2+^ signals seen by GCaMP3 imaging, we found that ASAP4.4-Kv imaging preserved fast temporal signal information, which GCaMP3 imaging failed to convey (Fig. 1d, e). ASAP4.4-Kv detected numerous signal changes associated with inflammation or nerve injury but GCaMP3 did not (Fig. 1e, f). In contrast, GCaMP3 Ca^2+^ signals reflected an increasing number of spontaneously activated neurons in the entire DRG after inflammation or nerve injury (Fig. 1g).

### ASAP4.4-Kv imaging permits visualization of mechanical stimuli (non-noxious to noxious)-evoked temporal summation of fast voltage signals

To understand how DRG neurons encode painful or non-painful mechanical stimuli, we applied stimulation of different strengths to the hindpaw, and visualized evoked ASAP4.4-Kv signals in DRG neurons. At low stimulation strength (light brush, 0.4 g, or 2 g von Frey; Fig. 2), small and transient subthreshold potential changes could be observed in mechanosensitive neurons (Fig. 2a), and only a few neurons exhibited hindpaw stimulation-evoked transient Ca^2+^ increases in naïve animals (Fig. 2b). However, peripheral inflammation or nerve injury led to a significant increase in membrane electrical signal summation, including both subthreshold and suprathreshold voltage signals (Fig. 2a, d), but not in Ca^2+^ responses (Fig. 2b, c, d). At an intermediate stimulation strength (100 g press), high-frequency voltage oscillations and dynamics were observed in neurons of naïve mice (Fig. 3a), while inflammation or nerve injury treatment produced exacerbated voltage fluctuations with larger amplitude and longer membrane depolarization (Fig. 3b, c, d). On the other hand, GCaMP3 Ca^2+^ imaging revealed increased population level activities in injured mice upon exposure to the same press stimulus (Fig. 3d, e). However, large variations in the magnitude of Ca^2+^ transients were found within the same DRG and across different treatment groups. Consequently, while the data were grouped, neither average amplitudes nor the mean AUCs of Ca^2+^ transients differed significantly between naïve or injured animals (Fig. 3d, g), despite the fact that increased activated cell numbers (Fig. 3d) and increased amplitudes of Ca^2+^ transients (Fig. 3f) were evident in some CFA-injured mice.

**Figure 2.**
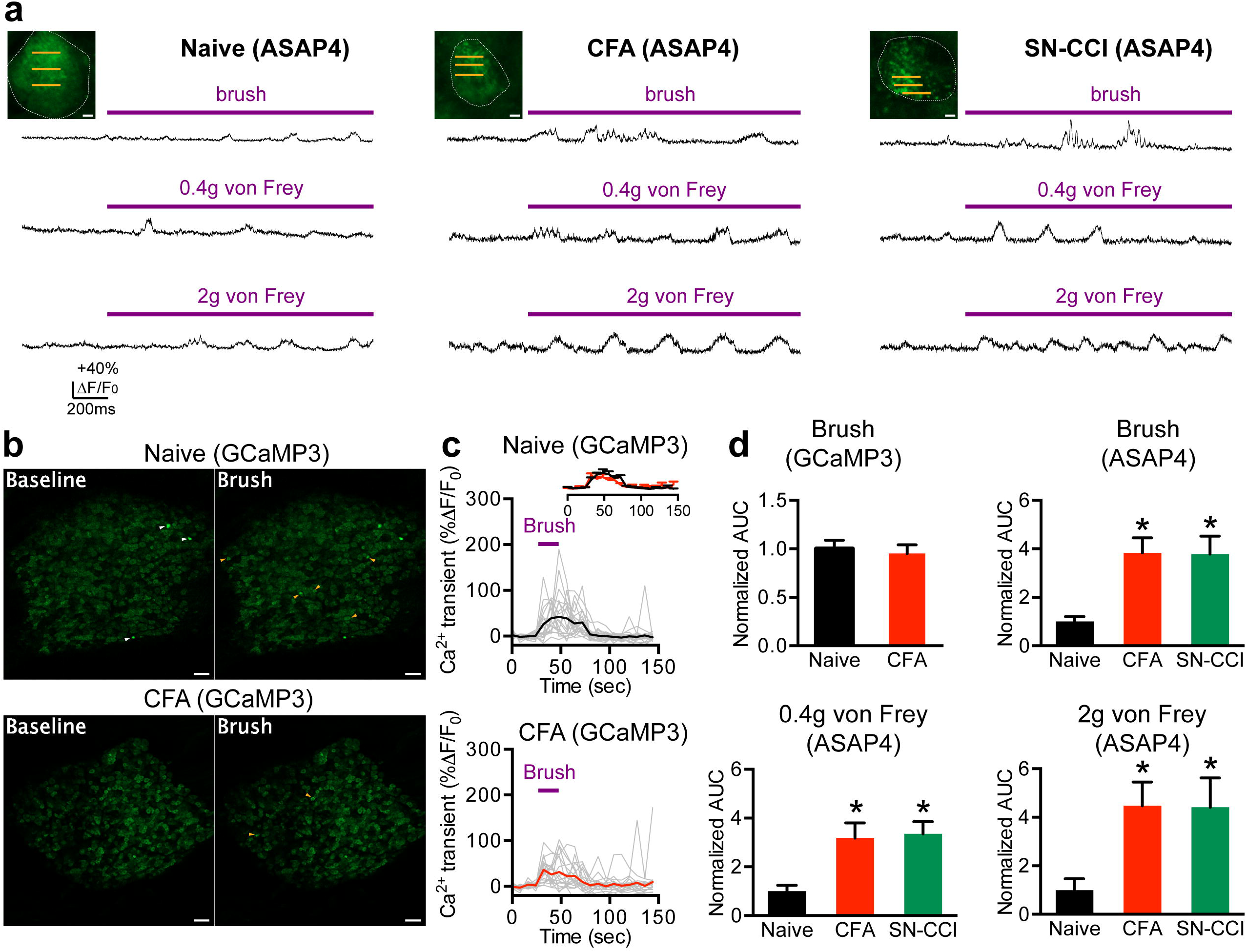
*In vivo* optical recording of mechanically (brush or von Frey)-induced neuronal activity in intact DRG neurons. (**a**) Optical voltage recordings of primary sensory neurons in response to the indicated stimuli in naïve, CFA, or SN-CCI animals. The traces under the image are different optical recordings from a single neuron. Purple bars indicate the timing of the stimulus application. Inset, images of DRG cell bodies expressing ASAP4.4. Yellow lines indicate 1.1 kHz line scan regions where ASAP4.4 fluorescent signals were acquired. Scale bar: 5 μm. (**b**) *In vivo* DRG Ca^2+^ imaging from naïve or CFA-treated Pirt-GCaMP3 mice. (*left*) Averaged images before brush stimulus was applied. (*right*) Averaged images after brush stimulus was applied. White arrowheads indicate spontaneously activated neurons without application of stimulus. Yellow arrowheads indicate DRG neurons activated by brush stimulus. Scale bar: 100 μm. (**c**) GCaMP3 Ca^2+^ transients from individual DRG neurons (grey traces) and averaged Ca^2+^ transients from naive or CFA animals (n=3 mice/treatment). (**d**) Mean area under the curve (AUC) of ASAP4.4 signals (n=3 mice/group; *P<0.05, Kruskal-Wallis test) and GCaMP3 Ca^2+^ transients (n=3 mice/group; P=0.7050, unpaired Student’s t-test) in stimulated L5 DRG neurons.

**Figure 3.**
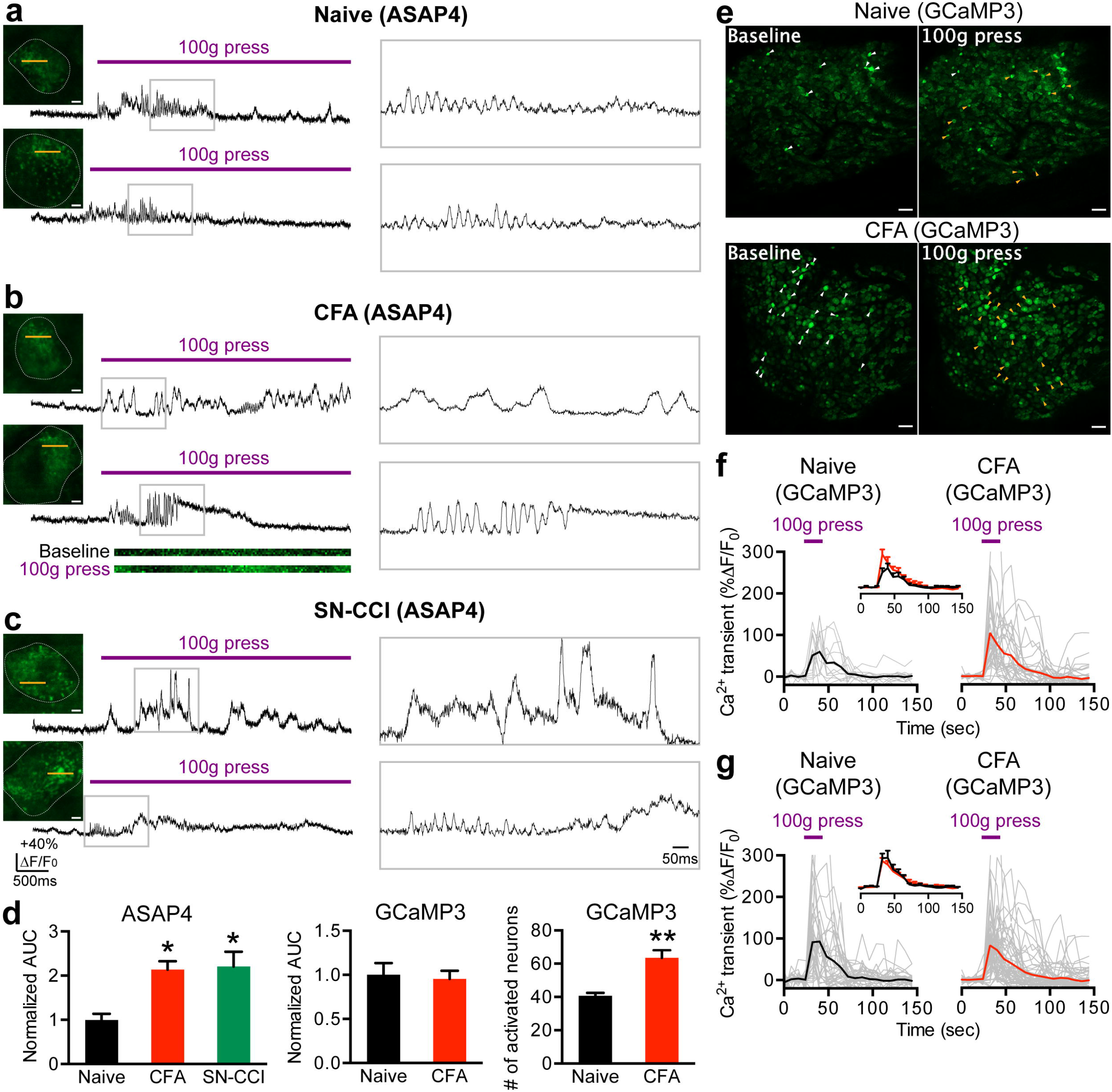
*In vivo* optical recording of mild press (100 g)-induced neuronal activity in intact DRG neurons. **(a-c)** Optical voltage recordings of primary sensory neurons in response to a single mechanical force (100 g) applied to the hindpaw of naïve **(a)**, CFA **(b),** or SN-CCI **(c)** animals. Insets, images of DRG cell bodies expressing ASAP4.4. Yellow lines indicate 1.1 kHz line scan regions where ASAP4.4 optical recording signals were acquired. Each trace is the response of a single DRG neuron; (*right*) expanded view of the boxed region. Purple bars indicate the timing of the stimulus application. Representative line scan images in (**b**) are shown under the trace. Scale bar: 5 μm. **(d)** Mean area under the curve (AUC) of ASAP4.4 signals (n=4 mice/treatment; *P<0.05, Kruskal-Wallis test) and GCaMP3 Ca^2+^ transients (n=3 mice/treatment; P=0.8942, Mann-Whitney U-test) in L5 DRG neurons in response to 100 g press, and the number of total activated neurons (n=5 mice/treatment; **P<0.01, Mann-Whitney U-test). **(e)** *In vivo* DRG Ca^2+^ imaging from naïve or CFA-treated Pirt-GCaMP3 mice. (*left*) Averaged images before application of mild press (100 g). (*right*) Averaged images after application of mild press (100 g). White arrowheads indicate spontaneously activated neurons in the absence of applied stimulus. Yellow arrowheads indicate DRG neurons activated by mild press. Scale bar: 100 μm. **(f, g)** Ca^2+^ transients from DRG neurons in response to mild press (100 g) applied to the hindpaw using Pirt-GCaMP3 mice (grey), from one **(f)** or three **(g)** animals for each treatment. Traces of averaged Ca^2+^ transients from each group are shown in black or red.

At the strongest mechanical stimulus (300 g), long-lasting membrane potential oscillations and fluctuations with sustained membrane depolarization were observed in DRG neurons of naïve mice (Fig, 4a), and voltage fluctuations in neuronal membranes were further aggravated by inflammation or nerve injury treatment (Fig. 4b, c). Similar to previous results, average Ca^2+^ transients differed only marginally between groups affected by inflammation or nerve injury (Fig. 4d, e, f), and increased numbers of activated cells were not evident (Fig. 4d). In addition, simultaneous *in vivo* dual color imaging of ASAP4.4-Kv (green) and mCyRFP3^27^, a cyan-excitable red fluorescent protein that can be used as a non-perturbing voltage-independent fluorescent marker as a control signal for ASAP4.4-Kv voltage imaging, demonstrated that the pattern of evoked electrical activity was distinguishable from rhythmic physiological motions arising from respiration or heartbeat (supplementary Fig. 4).

**Figure 4.**
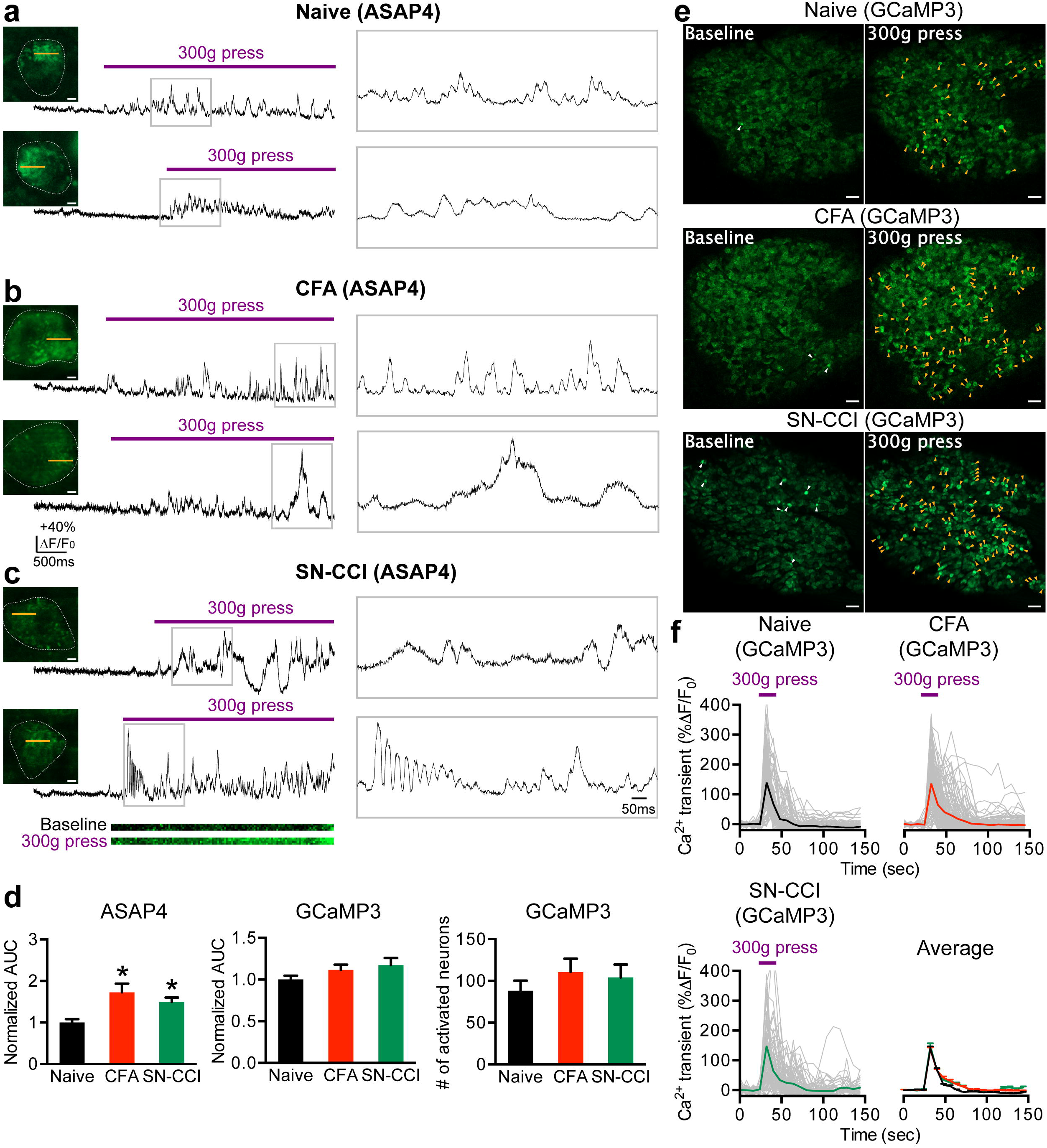
*In vivo* optical recording of strong press (300 g)-induced neuronal activity in intact DRG neurons. **(a-c)** Optical voltage recordings of primary sensory neurons in response to application of a single mechanical force (300 g) to the hindpaw of naïve **(a)**, CFA **(b),** or SN-CCI **(c)** animals. Insets, images of DRG cell bodies expressing ASAP4.4. Yellow lines indicate 1.1 kHz line scan regions where ASAP4.4 optical recording signals were acquired. Each trace is the response of a single DRG neuron; (*right*) expanded view of the boxed region shown on the right. Purple bars indicate the timing of the stimulus application. Representative line scan images in (**c**) are shown under the trace. Scale bar: 5 μm. **(d)** Mean area under the curve (AUC) of ASAP4.4 signals (n=4 mice/treatment; *P<0.05, Kruskal-Wallis test) and GCaMP3 Ca^2+^ transients (n=3 mice/treatment; P=0.8017, Kruskal-Wallis test) in L5 DRG neurons in response to 300 g press, and the number of total activated neurons (n=5 mice/treatment; P=0.5380, Kruskal-Wallis test). **(e)** *In vivo* DRG Ca^2+^ imaging from naïve, CFA, or SN-CCI using Pirt-GCaMP3 mice. (*left*) Averaged images before application of strong press (300 g). (*right*) Averaged images after application of strong press (300 g). White arrowheads indicate spontaneous Ca^2+^ activated neurons in the absence of applied stimulus. Yellow arrowheads indicate DRG neurons activated by strong press. Scale bar: 100 μm. **(f)** Ca^2+^ transients from DRG neurons in response to strong press (300 g) applied to the hindpaw of Pirt-GCaMP3 mice (grey) from all naïve, CFA, or SN-CCI animals. Traces of averaged Ca^2+^ transients from each group are shown in black, red, or green.

### ASAP4.4-Kv imaging reports thermal (heat or cold)-evoked voltage signals with high temporal fidelity

It has been reported that primary sensory neurons employ different strategies to encode heat vs. cold^28,29^. To discern how heat or cold is represented *in vivo*, we examined the ASAP4.4-Kv voltage signals from heat or cold-sensing neurons. In naïve mice, the membrane voltage dynamics during noxious heat (50°C) stimulation displayed a slowly depolarizing voltage ramp that returned to baseline within 300 ms (Fig. 5a). Noxious cold (0°C) stimulation, however, led to two distinctive forms of voltage activity: bursting or non-bursting (Fig. 6a). Bursting neurons displayed high frequency voltage oscillations and fluctuation behaviors, whereas non-bursting neurons exhibited slowly depolarizing voltage ramps similar to those seen in heat-sensing neurons (Fig. 6a). Inflammation or nerve injury, in turn, resulted in augmentation of membrane voltage fluctuation and electrical amplitude in both heat- and cold-sensing neurons (Fig. 5b, c, and Fig. 6b, c). Notably, stimulation by heat or cold was represented by distinct populational signals under various pain conditions.

**Figure 5.**
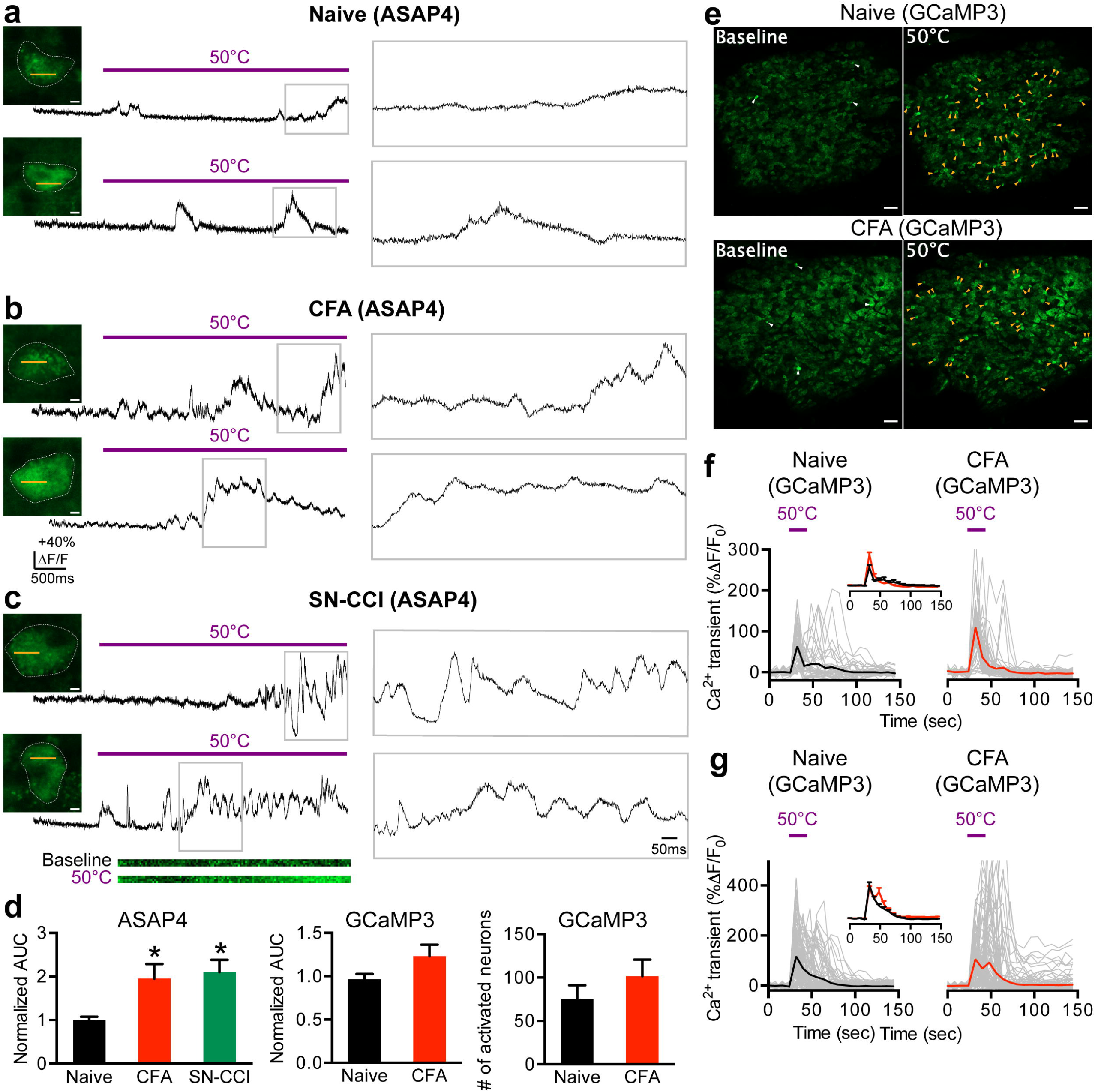
*In vivo* optical recording of neuronal activity in thermoreceptive neurons. **(a-c)** Optical voltage recordings of noxious heat (50°C)-sensitive primary sensory neurons from naïve (**a**), CFA (**b**), or SN-CCI (**c**) animals. Insets, images of DRG cell bodies expressing ASAP4.4. Yellow lines indicate 1.1 kHz line scan regions where ASAP4.4 optical recording signals were acquired. Each trace is the response of a single DRG neuron; (*right*) expanded view of the boxed region. Purple bars indicate the timing of the stimulus application. Representative line scan images in (**c**) are shown under the trace. Scale bar: 5 μm. **(d)** Mean area under the curve (AUC) of ASAP4.4 signals (n=4 mice/treatment; *P<0.05, Kruskal-Wallis test) and GCaMP3 Ca^2+^ transients (n=3 mice/treatment; P=0.3994, Mann-Whitney U-test) in L5 DRG neurons in response to noxious heat stimulus, and the number of total activated neurons (n=5 mice/treatment; P=0.3095, Mann-Whitney U-test). **(e)** *In vivo* DRG Ca^2+^ imaging from naïve or CFA-treated Pirt-GCaMP3 mice. (*left*) Averaged images before application of heat stimulus (50°C). (*right*) Averaged images after application of heat stimulus (50°C). White arrowheads indicate spontaneously activated neurons in the absence of applied stimulus. Yellow arrowheads indicate DRG neurons activated by heat (50°C). Scale bar: 100 μm. **(f, g)** Ca^2+^ transients from DRG neurons in response to heat (50°C) applied to the hindpaw of Pirt-GCaMP3 mice (grey), from one (**f**) or three (**g**) animals for each treatment. Traces of averaged Ca^2+^ transients of each group are shown in black or red.

**Figure 6.**
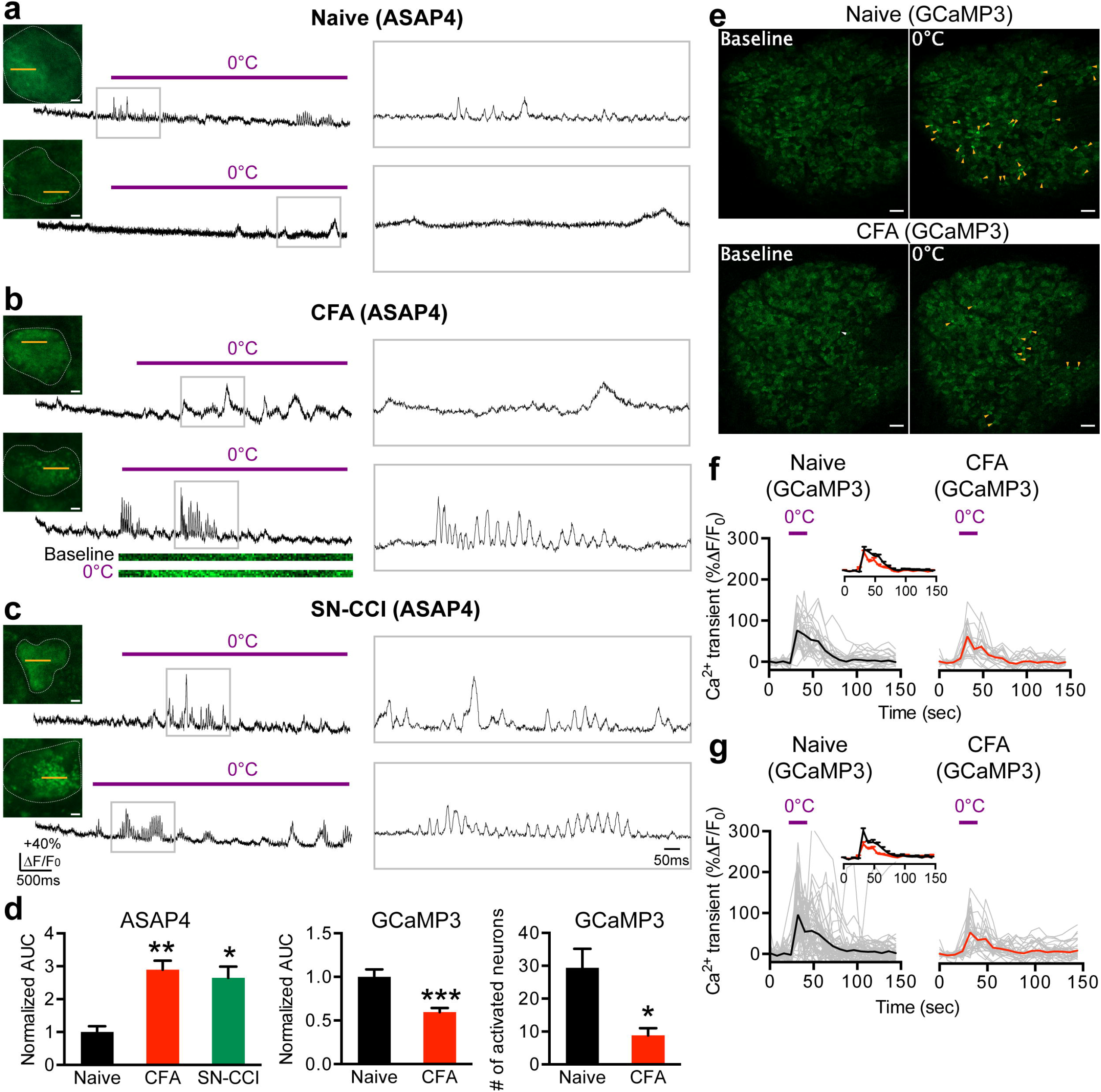
*In vivo* optical recording of neuronal activity in cold-sensing neurons. **(a-c)** Optical voltage recordings of cold (0°C)-sensitive primary sensory neurons from naïve (**a**), CFA (**b**), or SN-CCI (**c**) animals. Insets, images of DRG cell bodies expressing ASAP4.4. Yellow lines indicate 1.1 kHz line scan regions where ASAP4.4 optical recording signals were acquired. Each trace is the response of a single DRG neuron; (*right*) expanded view of the boxed region. Purple bars indicate the timing of the stimulus application. Representative line scan images in (**b**) are shown under the trace. Scale bar: 5 μm. **(d)** Mean area under the curve (AUC) of ASAP4.4 signals (n=5 mice/treatment; *P<0.05, **P<0.01, Kruskal-Wallis test) and GCaMP3 Ca^2+^ transients (n=3 mice/treatment; ***P<0.001, Mann-Whitney U-test) in L5 DRG neurons in response to noxious cold stimulus, and the number of total activated neurons (n=5 mice/treatment; P<0.05, Mann-Whitney U-test). **(e)** *In vivo* DRG Ca^2+^ imaging from naïve or CFA-treated Pirt-GCaMP3 mice. (*left*) Averaged images before application of cold stimulus (0°C). (*right*) Averaged images after application of cold stimulus (0°C). White arrowheads indicate spontaneously activated neurons without application of stimulus. Yellow arrowheads indicate DRG neurons activated by cold (0°C). Scale bar: 100 μm. **(f, g)** Ca^2+^ transients from DRG neurons in response to application of cold stimulus (0°C) to the hindpaw of Pirt-GCaMP3 mice (grey), from one (**f**) or three (**g**) animals for each treatment. Traces of averaged Ca^2+^ transients from each group are shown in black or red.

After CFA-induced inflammation, numerous DRG neurons were activated upon noxious heat stimulation (50°C) but numbers were similar between naïve and CFA groups (Fig. 5d, e), while fewer neurons displayed Ca^2+^ activity to noxious cold (0°C) compared to naïve animals (Fig. 6d, e). As with the previous mechanical stimuli, heat-induced increases in Ca^2+^ transients were observed in some DRGs of individual CFA-treated mice (Fig. 5f), but not in grouped DRGs (Fig. 5d, g). In contrast, cold-sensitive neurons displayed reduced Ca^2+^ transients after peripheral inflammation, both individually (Fig. 6f) and as a group (Fig. 6d, g). These results are consistent with previous reports that cold-mediated Ca^2+^ activity was lost in specific types of cold-sensing neurons following peripheral injury^29^. The discrepancy between voltage and Ca^2+^ signals in cold-sensing neurons suggests that, following peripheral inflammation, an individual sensory neuron still retains the ability to encode cold-specific sensory input; however, summation of the neuronal response to painful cold is suppressed by network activity in DRG.

### ASAP4.4 imaging reveals high potassium or capsaicin-evoked strong voltage fluctuations

Finally, we used the ASAP4.4-Kv voltage sensor to examine how DRG neurons encode noxious chemical nociception. In naïve mice, direct topical application of high potassium (50 mM KCl) or capsaicin (10 μM), a TRPV1 agonist which can initiate activity in nociceptive neurons, onto L5 DRG, resulted in an approximately 3-fold increase in voltage fluctuations over baseline (Fig. 7a, d). Both CFA and SN-CCI treatments significantly increased neuronal responses to KCl or capsaicin, with substantial increases in frequency and magnitude of dynamic membrane voltage fluctuations (Fig. 7b–d). When the same chemical treatments were performed on Pirt-GCaMP3 mice, we observed robust activation of a large population of DRG neurons within the DRG sensory ganglia (Fig. 7f, g). Topical application of capsaicin resulted in DRG neuronal activation primarily in the small and medium diameter neurons within all populations of DRG neurons imaged (Fig. 7g). Small and medium diameter neurons are nociceptors that typically express TRPV1 receptors. On average, the Ca^2+^ transients in activated neurons from injured mice were significantly higher than those from naïve animals (Fig. 7e). Compared to physical stimulation, direct chemical administration onto DRG neurons produced near-maximal Ca^2+^ transients and responses in most DRG neurons *in vivo*. These findings lead us to conclude that neuronal hypersensitivity is a common consequence of peripheral injury.

**Figure 7.**
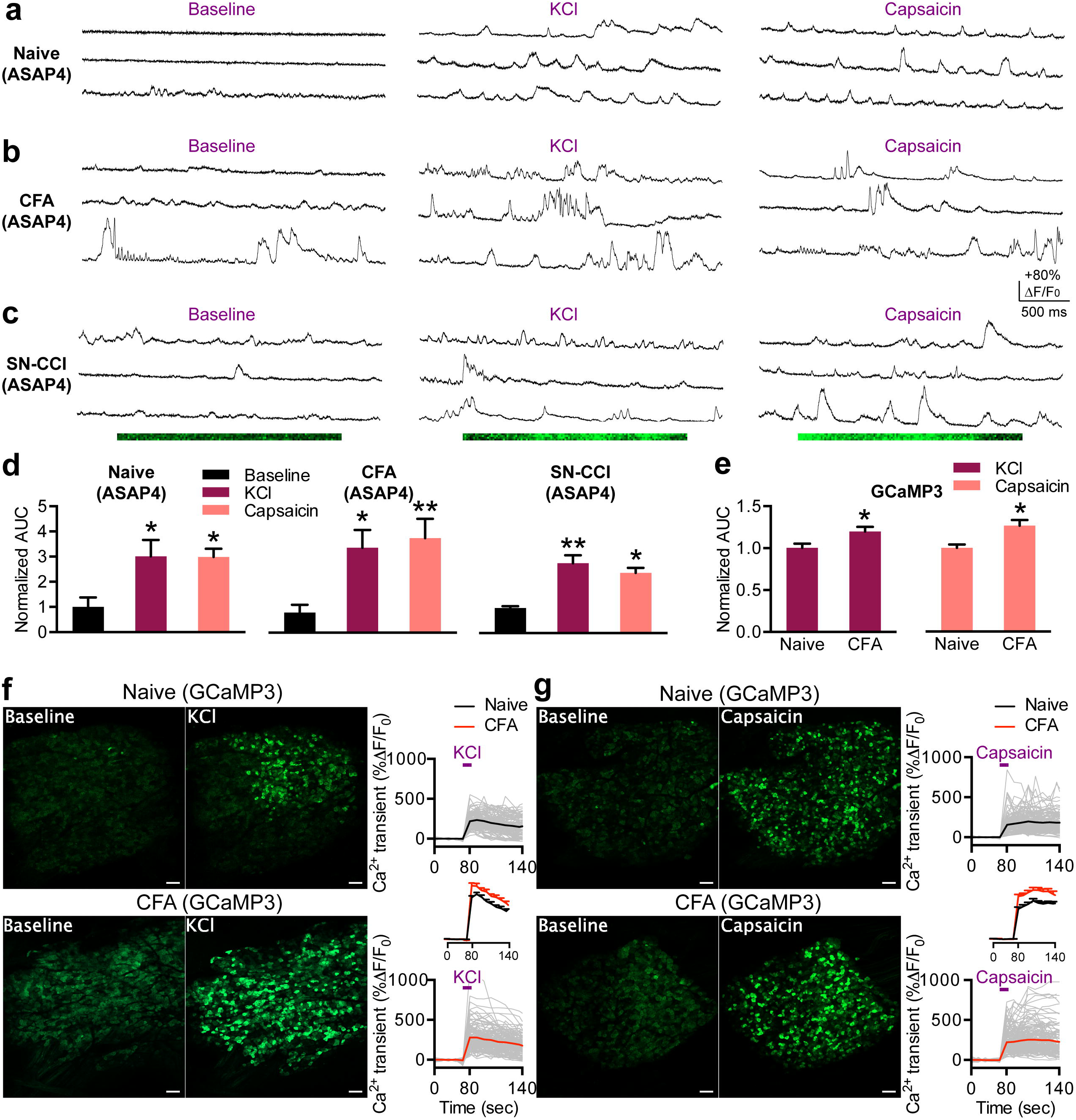
*In vivo* optical recording of intact DRG neurons in response to application of chemical stimuli (high potassium or capsaicin). **(a-c)** Representative traces of optical voltage recordings of primary sensory neurons before (baseline) and after topical application of KCl (50 mM) or capsaicin (10 μM) to L5 DRG from naïve **(a)**, CFA **(b),** or SN-CCI **(c)** animals. Representative line scan images in **(c)** are shown under the traces. **(d, e)** Mean area under the curve (AUC) of ASAP4.4 signals (**d**) (n=5 mice/treatment; *P<0.05, **P<0.01, Kruskal-Wallis test) and GCaMP3 Ca^2+^ transients (**e**) (n=3 mice/treatment; *P<0.05, Mann-Whitney U-test) in L5 DRG neurons in response to indicated chemical stimuli. **(f, g)** *In vivo* DRG Ca^2+^ imaging from naïve or CFA-treated Pirt-GCaMP3 mice. (*left*) Averaged images before application of indicated chemical stimuli. (*middle*) Averaged images after application of indicated chemical stimuli. Scale bar: 100 μm. (*right*) Ca^2+^ transients (gray traces) from DRG neurons in response to topical application of KCl (**f**) or capsaicin (**g**) onto DRG neurons from naïve or CFA-treated Pirt-GCaMP3 mice. Traces of averaged Ca^2+^ transients from each group are shown in black or red.

## Discussion

GEVIs have been successfully used in analysis of brain regions *in vivo* in awake behaving mice^15,16,30^, and have encouraged neuroscientists to explore and unlock the full potential of the technological advances. Our current study reports the use of an improved ASAP-family GEVI, ASAP4.4-Kv, to track both spontaneous and evoked voltage activity of mouse primary sensory neurons *in vivo*. The ASAP4.4-Kv voltage sensor allows direct visualization of distinct temporal features of neuronal dynamics, subcellular voltage dynamics, plasticity induction, and neuronal coding in DRG, the analysis of which have been largely inaccessible and technically challenging in live animals. The ASAP4.4-Kv voltage sensor provides the means for understanding how primary sensory neurons, especially DRG neurons, function or fail to function (changes in dynamics, status, and/or pattern) at any given time under physiological and pathological conditions.

### GEVI imaging as a powerful tool complementary to GECI imaging

ASAP3-Kv, a previous ASAP family GEVI with desirable responsivity and SNR for *in vivo* use, has a negative slope relationship between voltage and fluorescence^15,16^. But, new ASAP4.4-Kv produces a depolarization-dependent increase in fluorescence intensity^17^, as do most GCaMP GECIs^4,26^. This new feature, combined with somatic targeting and optimal brightness, greatly improves the utility of the GEVI ASAP4.4-Kv for *in vivo* recording of neuronal electrical signals and activity. In our *in vivo* DRG voltage sensor imaging studies, we could visualize sparse signals from the membrane surface of neuronal soma expressing fluorescent ASAP4.4-Kv with imaging depths of <20 μm below the meninges membrane. We demonstrated superiority of *in vivo* GEVI recording for resolving long-standing debates over the temporal attributes of neuronal coding by direct comparison with *in vivo* GECI imaging modalities previously developed by our lab^4^. Compared to GECIs, the main advantage of GEVIs is their ability to examine nonspiking (subthreshold) and/or spiking (action potential) electrical activity, because subthreshold membrane potential fluctuation and oscillation do not greatly affect internal Ca^2+^ levels or dynamics. ASAP4.4-Kv is able to identify non-spiking subthreshold voltage fluctuation events in DRG neurons with duration times in the millisecond range, an order of magnitude faster than the signal integration time of the Ca^2+^ indicator GCaMP3 or other advanced GCaMPs.

Unexpectedly, spontaneous neuronal dynamics revealed by ASAP4.4-Kv exhibited cell-to-cell coupled synchronous electrical events following injury that were normally indiscernible in GCaMP3 imaging across all DRG neurons. Highlighting the importance of *in vivo* voltage imaging, ASAP4.4-Kv imaging fully unmasked altered DRG neuronal electrical activities and signals resulting from peripheral injury at the single-cell level in their native environment. In contrast, variable Ca^2+^ activities could be observed in *in vivo* Prit-GCaMP3 Ca^2+^ imaging, but significant effects were diminished in comparisons of larger numbers of DRG neurons from multiple animals. Another fascinating aspect of fluorescence voltage sensors is their ability to map not only subthreshold depolarizing (excitatory) inputs but also hyperpolarizing (inhibitory) events that occur constantly in almost all neurons^31^. Previous studies have used voltage indicators to monitor membrane hyperpolarization in cultured neurons^32^, brain slices^33^, or in freely moving mice^34^. The hyperpolarizing voltage signals detected by ASAP4.4-Kv in our DRG recordings were evident in response to strong stimuli, and were detected as fluorescence intensity dropped to levels below the pre-stimulation baseline. In contrast, GECI imaging can only detect excitatory inputs, and lacks ability to detect signals related to inhibitory inputs. Thus, a plausible use of GEVI imaging would be to examine possible inhibitory signaling involved in controlling peripheral nociceptive or nonnociceptive transmission, or to examine excitatory or/and inhibitory signal summation and integration within sensory ganglia.

Given the heterogeneity of sensory neurons, integrated signals from large-scale DRG neurons need to be collected at high spatiotemporal resolution to establish cell-type or modality-specific coding strategies. The intrinsic slow kinetics of GECIs permits mapping large numbers of neuronal assemblies in their native environments with conventional confocal microscopic approaches. Intensive studies using GECIs have characterized sensory coding of heat or cold^28,29,35–37^, mechanical^35,37,38^, or chemical stimuli^39^ in health and disease conditions. Due to the intrinsic fast kinetics of GEVIs, simultaneous imaging of dozens, hundreds, or thousands of DRG neurons at high spatial (millimeters) and temporal (milliseconds) resolution is an extremely challenging task and limited by current technological advances. To overcome these limitations, we adapted a conventional, low cost, strong laser intensity, upright laser-scanning confocal microscope to be used as a versatile platform, allowing optical reporting of dynamic neuronal activity in DRG neurons at high spatial and temporal resolution by combining GECI-based Ca^2+^ signals and GEVI-based voltage signals. With the continuous advances in voltage indicators and optical instruments, simultaneous voltage recording of enormous numbers of neurons in live intact peripheral tissues will enable dissecting functional connectivity in DRG circuits and mapping neuronal coding strategies with better high-throughput and greater accuracy.

### Future outlook

We previously described the phenomenon of coupled neuronal activation within DRG in mouse models of inflammatory or neuropathic pain^4^. Here, our *in vivo* voltage imaging results provide evidence of electrically synchronous neuronal activity between DRG neurons, further supporting increased neuronal ‘cross talk’ as an underlying mechanism in the development of hyperalgesia and allodynia. In our ongoing work, we aim to elucidate the pathway and mechanism of neuron-to-neuron transmission.

Our current implementation combining both GEVI and GECI imaging should allow detailed investigations of the relationship between suprathreshold somatic voltage signals and the corresponding Ca^2+^ dynamics at single-cell-, population-, or modality-specific levels. Simultaneous sub-millisecond voltage and Ca^2+^ imaging using a voltage-sensitive dye and GECI has been performed on Purkinje neurons in awake animals, and has demonstrated high spatiotemporal variations of suprathreshold voltage signals and Ca^2+^ transients between dendritic segments^40^. For *in vivo* primary sensory neuron studies, dual GEVI and GECI neuronal labeling with different fluorescence spectra will help in the analysis of correlation or integration between suprathreshold voltage signals and the resulting Ca^2+^ transients at high spatiotemporal resolution.

### Limitations of this study

As a latest ASAP-family sensor, positively tuned ASAP4.4-Kv with optimal fluorescence response and SNR enables identifying and tracking each suprathreshold voltage spike during the designated time in our *in vivo* DRG recording. In two-photon optical imaging in deep layers of *in vivo* brain, a previously used ASAP-family GEVI, ASAP3, reliably detected single spikes and resolved spikes in bursts, with appreciable optical spike amplitudes^15,16^. However, in our *in vivo* DRG studies, a few spikes were detectable from neurons expressing ASAP4.4-Kv within a short period of time. Typically, minor suprathreshold voltage spikelets were visible at the peaks during the periods of voltage oscillation. This may be due to different physiological firing patterns in peripheral sensory neurons, or relatively less sensitivity compared to ASAP3, or a combination of these.

Another limitation of our study was the use of viruses to transduce the ASAP4.4-Kv gene into DRG neurons. Viral vectors can be used in a variety of species and for cell type-specific expression, while gene delivery via virus injection can lead to uneven expression strength related to site of injection, and a limited time window of acceptable expression level^5,41,42^. Transgenic animals should overcome these limitations as demonstrated by our studies using the *Pirt* promotor to selectively drive strong GCaMP3 expression in DRG neurons^4^. Currently, transgenic mouse lines expressing the GEVI have been reported for *in vivo* studies in olfactory cells^43^. Future engineering efforts could focus on transgenic GEVI mouse lines for selective expression in targeted tissues, which promise spatially homogeneous transgene expression and long-term time windows for GEVI imaging.

In conclusion, ASAP4.4-Kv voltage imaging opens new avenues to explore the basic principles of DRG neuron coding, and of the cellular basis for perceptual changes in somatosensation by providing high temporal resolution of individual neurons. The combination of GEVI and GECI imaging allows a more temporally and spatially precise characterization of the neuronal coding and integration strategies in the peripheral somatosensory system.

## Methods

### Animal models

All experiments were performed in accordance with a protocol approved by the Institutional Animal Care and Use Committee at University of Texas Health Science Center at San Antonio (UTHSA). C57BL/6J mice (body weight 20–30 g) were obtained and bred in-house. Animals were group housed unless otherwise noted, provided with food and water ad libitum, and kept on a 14/10 light/dark cycle at 23°C. To generate CFA inflammatory injury mice, we made a 1:1 mixture of complete Freund’s adjuvant (CFA): saline, and injected 50 μL subcutaneously into the glabrous skin of the hindpaw. *In vivo* imaging was performed 1-3 days following CFA injection. To generate sciatic nerve (SN) chronic constriction injury^23^, mice were anesthetized by intraperitoneal (i.p.) injection of ketamine/xylazine (0.1/0.015 mg/g body weight). SN was exposed mid-thigh by a small incision and separated from surrounding tissue. Ligatures were loosely tied using 3-0 silk thread around SN. The incision was closed using sutures, and mice were used for *in vivo* imaging 7–10 days later.

Pirt-GCaMP3 mice were generated and described as done in a previous study^4,26^. Briefly, transgenic animals were generated by targeted homologous recombination to replace the entire coding region of the *Pirt* gene with the GCaMP3 sequence in frame with the *Pirt* promoter.

### ASAP4.4-Kv virus delivery

AAV8-hSyn-ASAP4.4-Kv and AAV8-hSyn-ASAP4.4-Kv-mCyRFP3.WPRE^17^ were generated by the Stanford Viral Core. For the experiments in cell culture, male or female 2 to 3 weeks-old mice were used for intrathecal delivery of the virus to DRG neurons. For *in vivo* imaging, 2 to 4 months old mice from both sexes were used for intrathecal delivery of virus to peripheral neurons. For intrathecal delivery, mice from both sexes were first anesthetized with isoflurane, shaved and disinfected. The AAVs were diluted in sterile, isotonic saline. A volume of 30 μl containing 2×10^12^ virus particles/ml was injected intrathecally (i.t.) by direct lumbar puncture using a 28½-gauge needle and insulin syringe (Becton Dickinson, Franklin Lakes, NJ). A reflexive flick of the tail indicated proper needle entry location for intrathecal injection. Following the injection, the animals were returned to recovery cages where they remained for 5–7 weeks until imagining or electrophysiology experiments were performed.

### DRG exposure surgery

DRG exposure surgery was carried out as previously described^4^. Briefly, mice were anesthetized with i.p. injection of ketamine/xylazine (0.1/0.015 mg/g body weight). Mice were kept on a heating pad to maintain body temperature at 37±0.5°C, which was monitored by a rectal probe. Their backs were shaved, and ophthalmic ointment was applied to their eyes to prevent drying. The transverse processes of lumbar L5 were exposed, and the surface aspect of the bone covering the DRG was carefully removed to expose the underlying DRG without damaging the DRG or spinal cord. Bleeding was gently stopped using styptic cotton or gel foam.

### *In vivo* imaging

For *in vivo* imaging of the whole L5 DRG, mice were placed on a custom-built tilted stage and their spines were secured with custom-built clamps to minimize movement due to breathing and heartbeat. The stage was affixed under a LSM 800 confocal laser-scanning microscope (Carl Zeiss, Inc) equipped with upright 5×, 10×, and 40× objectives. The isoflurane-anesthetized animals (1%–2%, vol/vol in 100% O_2_) were maintained at 37±0.5°C by a heating pad during the imaging process. Z-stack imaging, which can cover the entire L5 DRG, was typically acquired at eight to ten frames using a 10× C Epiplan-Apochromat objective (0.4-NA, 5.4-mm free working distance, Carl Zeiss) at typically 512×512 pixel resolution with lasers tuned at 488 nm and at 561 nm and emission at 500–550 nm for green and 620–700 nm for red fluorescence. DRG neurons were at the focal plane, and imaging was monitored during the activation of DRG neuron cell bodies by peripheral hindpaw stimuli.

For the recording of ASAP4.4-Kv fluorescent signals, z-stack imaging was first performed to localize the individual spiking neurons which responded to peripheral hindpaw sensory stimuli. Next, bidirectional line scanning along neuronal cell bodies was performed on the chosen cell to achieve fast ASAP4.4 imaging at 1.1 kHz. After adjusting focus depths to avoid artifacts of membrane motion, around 7700 lines (7 s) at <1 ms per line, 128 × 4 pixels in image size, and 0.12 μm in pixel size were acquired per stimulus. For reporting cell-to-cell communication, around 5500 lines (5 s) at <1 ms per line were imaged at 1024 × 1 pixels in image size and 0.02 μm in pixel size. Fluorescent signals of each line were integrated to produce a movie file of fluorescent trace over time. For ASAP4.4 signal analysis, the first 500–2000 lines were typically discarded due to photobleaching, and fluorescent traces with strong motion artifacts were also excluded in the analysis.

### Stimulus delivery during imaging experiments

Mechanical or thermal stimuli were applied on the ipsilateral hindpaw in the following order: brush, 0.4 g von Frey, 2 g von Frey, 100 g press, 300 g press, heat (50°C), or cold (0°C). Paw stimuli with 100 g or 300 g press force were delivered using a rodent pincher analgesia meter, and press force was controlled manually by the experimenter. For the ASAP4.4 imaging, the duration of the external stimuli was 4–5 s after 2–3 s of baseline imaging. For the GCaMP3 imaging, a time series of a total of 20 cycles was performed for each stimulus. The first 5 cycles (40–50 s) of z-stack images were captured for baseline activity determination, and images for another 5 cycles (40–50 s) were taken when paw or DRG stimuli were applied. Images were continuously taken for a total of 20 cycles. At the end of experiments, 20 μl of 50 mM KCl or 10 μM capsaicin was applied dropwise to the L5 DRG following 5 cycles of baseline imaging. After an incubation period of 1–2 cycles (5–10 s), KCl or capsaicin solution was removed by Kimwipe tissue, and then an additional 5 cycles of images were captured. Each stimulus was separated by an interval of 3–10 mins resting time for mice to avoid sensitization of neurons.

### DRG culture

ASAP4.4-transduced mice of either sex (6–7 weeks old, 4 weeks after intrathecal delivery of AAV8-hSyn-ASAP4.4-Kv virus) were anesthetized with isoflurane, and euthanized by decapitation. DRG were removed bilaterally at L3–L5 and incubated in collagenase (Worthington) and dispase (Sigma-Aldrich) for 40 min at 37°C with gentle agitation every 10 min. The dissected DRG neurons were then triturated, centrifuged, and resuspended in Dulbecco’s Modified Eagle Medium (DMEM, Gibco, Grand Island, NY) supplemented with 10% fetal bovine serum (FBS, Gibco), 100 ng/ml nerve growth factor (NGF, Harlan, Indianapolis, IN), 1% penicillin/streptomycin (Gibco), and 1% L-glutamine (Gibco), and then placed on coverslips coated with poly-D-lysine and laminin (Corning, Corning, NY). Cultures were maintained at 37°C, 5% CO_2_ for 24 hr prior to electrophysiologic recordings.

### *In vitro* electrophysiologic recording and green fluorescence imaging with ASAP4.4

Whole-cell patch-clamp recordings were performed under voltage-clamp mode using an Muticlamp 700B amplifier and pClamp11 software (Molecular Devices), and data were digitized using an Axon Instruments Digitizer. Pipette membrane capacitance was compensated, and currents were sampled at 10 kHz. Glass pipettes (3-4 MΩ resistance, World Precision Instruments (Sarasota, FL)) were filled with an intracellular solution containing the following: 140 mM K-gluconate, 5 mM KCl, 10 mM HEPES, 5 mM MgCl_2_, 4 mM Mg-ATP, 0.3 mM Na-GTP, pH 7.2. The coverslips containing DRG neurons were transferred to the recording chamber and continuously perfused with recording solution containing: 125 mM NaCl, 2.5 mM KCl, 2 mM CaCl_2_, 1 mM MgCl_2_, 1.5 mM NaH_2_PO_4_, 15 mM NaHCO_3_ and 10 mM D-glucose (pH 7.4), and bubbled with 5% CO_2_/95% O_2_.

Fluorescence traces were acquired with cells using whole-cell voltage-clamp mode. Step voltage was applied to change the membrane potential from a holding voltage of −70 mV to command voltages at −100, −40, +30, or +100 mV in a series of subsequent steps for 0.5–1 s. ASAP4.4-Kv expressing DRG neurons were imaged on an upright Zeiss Examiner.A1 microscope fitted with a 40× water-immersion objective (0.75-NA, 2.1-mm free working distance, Carl Zeiss) and with an Axiocam 705 color camera (Carl Zeiss). Images were sampled at 5 Hz.

### *In vivo* imaging data analysis

To analyze confocal line-scan imaging of ASAP4.4, fluorescence imaging data were extracted from raw image data, and time-dependent fluorescence traces for each neuron were revealed using Mean ROI function in Zen blue software. Because presentation of peripheral stimuli evoked spatially differentiated, large optical signals that were distinguishable from the stimulus-independent component, we averaged the first 1–2 s before the stimulus onset and designated that as the baseline fluorescence (F_0_). Baseline-normalized amplitudes in the region of interest (ROI) over time were expressed as (F–F_0_)/F_0_×100% against time. Some experiments were excluded if rundown exceeded 30%.

For GCaMP3 imaging data analysis, individual responding neurons were verified by visual examination and confirmed when the fluorescent intensity of ROI during stimulus was 15% higher than baseline signals using the Mean ROI function in Zen blue software. Time series recorded fluorescence changes were exported to Excel, and were analyzed using GraphPad prism. The average fluorescence intensity in the baseline period was taken as F_0_, and was measured as the average pixel intensity during the first two to five frames of each imaging experiment. Relative change in fluorescence intensity was measured using the formula ΔF/F_0_ (%) = (F–F_0_)/F_0_×100%.

### Statistical methods

Group data were expressed as mean ± standard error of the mean (S.E.M.). Student’s unpaired t-tests, Mann-Whitney U-tests, one-way ANOVA with a post-hoc Dunnett’s t-test, or Kruskal-Wallis test, as appropriate, were employed for comparisons; *P<0.05, **P<0.01, ***P<0.001, ****P<0.0001. All statistical tests are indicated in figure legends.

## Acknowledgements

This study was funded by the National Institutes of Health Grant (R01DE026677 to Y. S. K), UTHSA Startup (Y. S. K), and a Rising STAR Award (Y. S. K) from University of Texas system.

## Author Contributions

Y.Z. and Y.S.K contributed to study design with assistance from J.S., R.G., H.S., H.I., and M.L. M.C. and M.L. developed the predecessor to ASAP4.4 voltage sensor. D.S. finalized and made the ASAP4.4 sensor. G.Z. cloned the ASAP4.4 viral constructs. M.C. made the ASAP4.4 virus. Y.S.K. contributed to data interpretation and manuscript revision. Y.Z. conceived the project and performed all experiments except where noted, and drafted the paper. R.G. and J.S. maintained, set up mating, took care of mice, and performed genotyping. R.G. assisted with GCaMP imaging work. Y.S.K. supervised all aspects of the project and wrote the paper.

---

**Supplementary Figure 1. Representative images of *in vivo* entire L5 DRG neurons expressing ASAP4.4. (a, b**) Distribution of ASAP4.4-expressing neurons (green) in L5 DRGs and ASAP4.4-expressing L5 DRG neurons that project to hindpaw (yellow from green/red merged). Retrograde tracer WGA- **(a)** or Dil-labeled **(b)** somata in L5 DRG. WGA (10 μl of 1 mg/mL, s.c. plantar surface) or Dil (10 μl of 1 mg/mL, s.c. plantar surface) was injected into a hindpaw and imaged 7 days later. **(c)** Effects of CFA or SN-CCI treatments on ASAP4.4 fluorescence level in L5 DRG. White arrowhead points to single DRG neurons which were selected for *in vivo* single-cell voltage imaging. Scale bar: 100 μm.

**Supplementary Figure 2. ASAP4.4 is well suited for optical imaging responses across the physiological voltage range in DRG neurons. (a)** ASAP4.4 responses in a dissociated DRG neuron to voltage steps from −70 mV holding potential to −100 mV or to +100 mV. Yellow arrowhead indicates a neuron that was voltage clamped to report ASAP4.4 fluorescence changes. White arrow indicates another neuron on the same coverslip which did not show a significant change in ASAP4.4 fluorescence. Scale bar: 10 μm. **(b)** Mean ASAP4.4 response to different command voltages (−100 mV to +100 mV) in DRG neurons (n=3 cells). Responses were normalized to fluorescence at the −70 mV holding potential.

**Supplementary Figure 3. Adjacent cell-to-cell electrical synchronization is attenuated by the gap junction blocker, carbenoxolone (CBX). (a)** Representative ASAP4.4-Kv optical recordings showing cell-to-cell electrical synchronization between a pair of adjacent DRG neurons in a CFA animal. Yellow line indicates 1.1 kHz line scan regions where ASAP4.4 fluorescent signals were acquired. Scale bar: 10 μm. **(b)** Electrical de-synchronization between a pair of adjacent DRG neurons approximately 1 hr after systemic gap junction blocker, CBX, injection (100 mg/kg, i.p.). The subthreshold voltage dynamics from recording neurons were gradually diminished following de-synchronization by gap junction blocker. **(c)** Quantification of synchronous pairs in control and after systemic CBX injection (*P<0.05, Mann-Whitney U-test). **(d)** Correlation coefficient was calculated from DRG neuron pairs in control and after systemic CBX injection (*P<0.05, Mann-Whitney U-test). Neuron pairs exhibit electrical correlation, whereas the correlation is attenuated by gap junction blocker. Fluorescence traces showing respiratory motion or heartbeat-like rhythmic events, or digital artifacts were not considered as electrical synchronization in the analysis.

**Supplementary Figure 4. ASAP4.4 optical signal recording is not affected by motion. (a)** Fusion of ASAP4.4 to a non-perturbing voltage-independent marker, mCyRFP3, shows sparse membrane expression and tight correspondence between ASAP4.4 (green signal) and mCyRFP3 (red signal). Scale bar: 5 μm. **(b)** Representative ASAP4.4 (green) and mCyRFP3 fluorescence in response to a single application of mechanical force (300 g) to the hindpaw of a naïve mouse. Note that bleed-through from the green channel was seen in the red channel fluorescence, but rhythmic respiratory motions or heartbeats in the live animals did not interfere with recording the ASAP4.4 optical signal.

## Supplemental Movies

**Supplementary Movie 1, related to Figure 4. Representative *in vivo* optical ASAP4.4 recording of a L5 DRG neuron from SN-CCI mouse in response to 300g press applied to hindpaw.** Each frame takes 0.91 ms, 1.1 kHz speed. Note that the movie resolution became reduced by conversion from Zen blue (Zeiss) software to movie format AVI file.

**Supplementary Movie 2, related to Figure 5. Representative noxious heat (50°C)-induced dynamic voltage imaging of a L5 DRG neuron from SN-CCI mouse.**

**Supplementary Movie 3, related to Figure 6. Representative noxious cold (0°C)-induced dynamic voltage imaging of a L5 DRG neuron from CFA-injected mouse.**

**Supplementary Movie 4, related to Figure 7. Representative *in vivo* optical ASAP4.4 recording of a L5 DRG neuron from SN-CCI mouse before application of chemical stimuli onto DRG neuron.**

**Supplementary Movie 5, related to Figure 7. Representative *in vivo* optical ASAP4.4 recording of a L5 DRG neuron from SN-CCI mouse in response to topical application of KCl (50 mM).**

**Supplementary Movie 6, related to Figure 7. Representative *in vivo* optical ASAP4.4 recording of a L5 DRG neuron from SN-CCI mouse in response to topical application of capsaicin (10 μM).**

**Supplementary Movie 7, related to Figure 3. Representative *in vivo* entire L5 DRG Pirt-GCaMP3 Ca^2+^ imaging from naïve and CFA-treated mice in response to 100 g mechanical press to hindpaw.**

**Supplementary Movie 8, related to Figure 5. Representative *in vivo* entire L5 DRG Pirt-GCaMP3 Ca^2+^ imaging from naïve and CFA-treated mice in response to noxious heat (50°C) stimulus to hindpaw.**

