## Supplementary figures and images for "Imaging sensory transmission and neuronal plasticity in primary sensory neurons with genetically-encoded voltage indicator, ASAP4.4-Kv"

### Supplemental Figure1

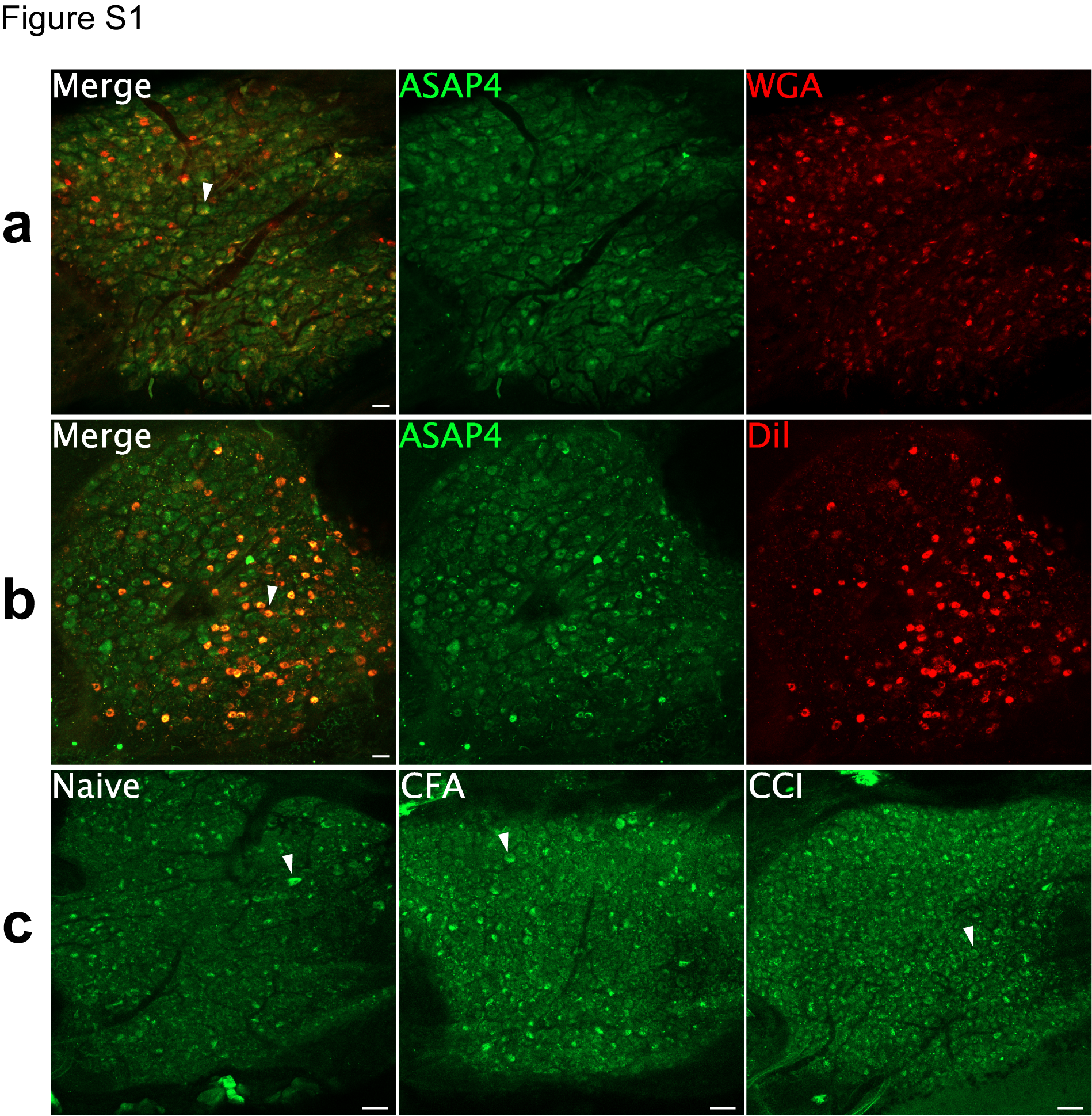

### Supplemental Figure2

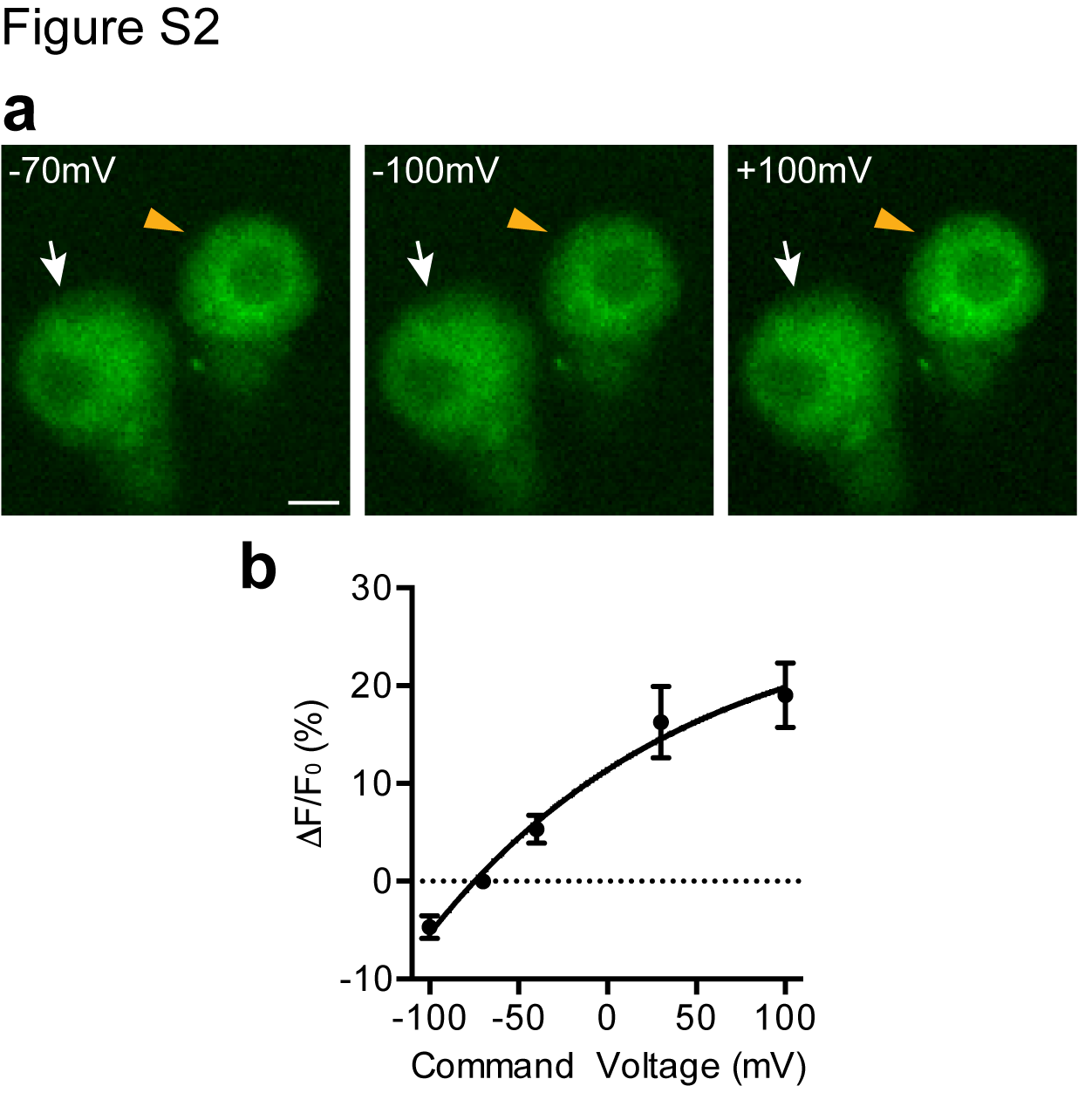

### Supplemental Figure3

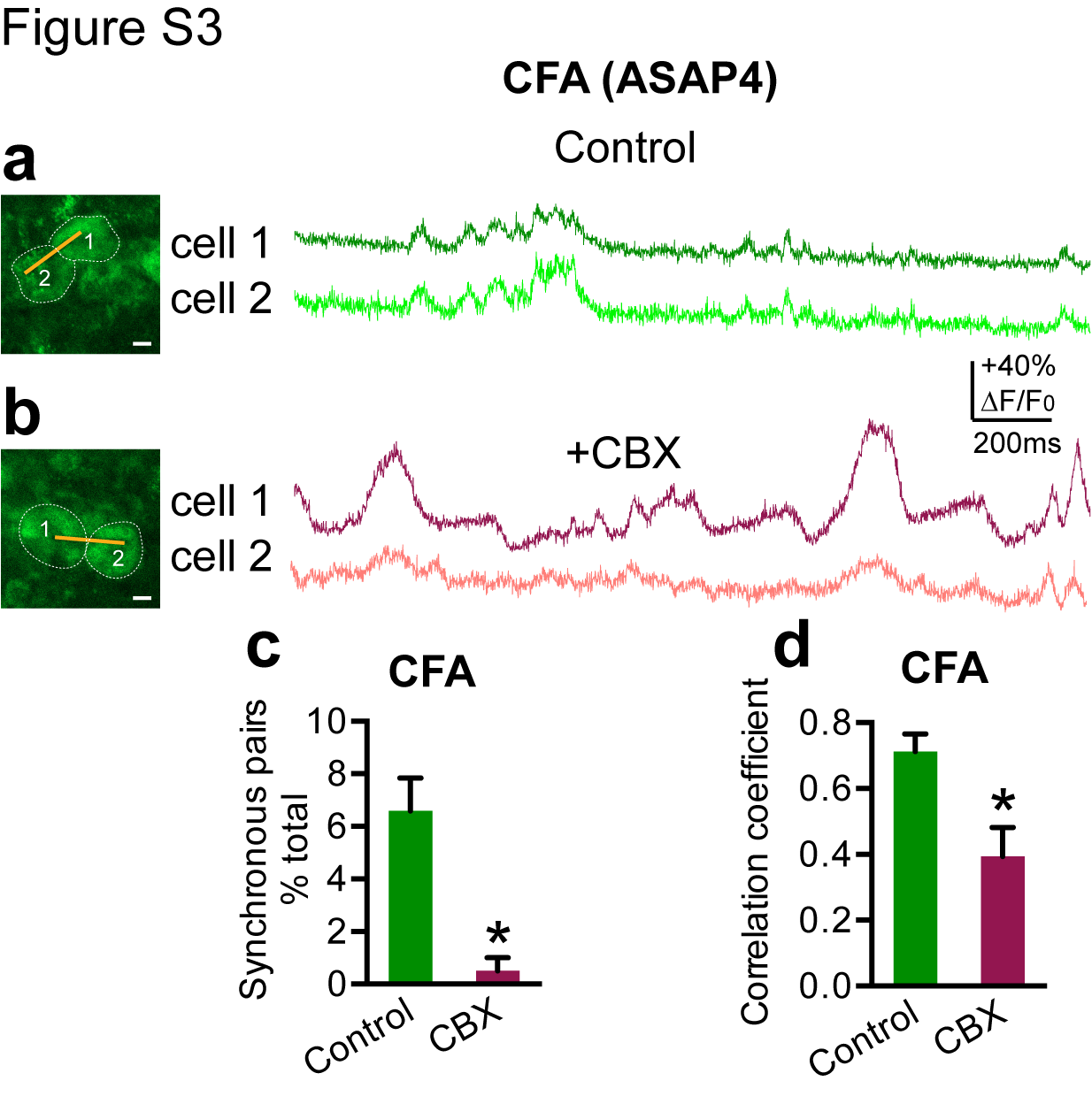

### Supplemental Figure4

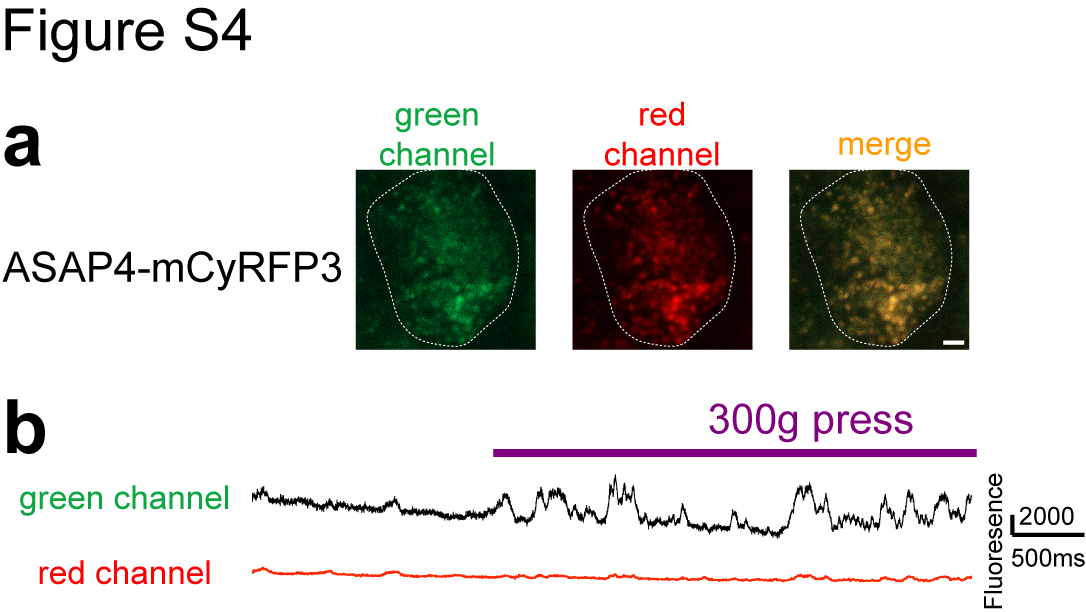
